# Dynamic Expedition of Leading Mutations in SARS-CoV-2 Spike Glycoproteins

**DOI:** 10.1101/2021.12.29.474427

**Authors:** Muhammad Hasan, Zhouyi He, Mengqi Jia, Alvin C. F. Leung, Kathiresan Natarajan, Wentao Xu, Shanqi Yap, Feng Zhou, Shihong Chen, Hailei Su, Kaicheng Zhu, Haibin Su

**Author notes:** These authors contribute equally (Muhammad Hasan, Zhouyi He, Mengqi Jia, Alvin C. F. Leung, Kathiresan Natarajan).

## Abstract

Throughout the coronavirus disease 2019 (COVID-19) pandemic, the continuous genomic evolution of its etiological agent, severe acute respiratory syndrome coronavirus 2 (SARS-CoV-2), has generated many new variants with enhanced transmissibility and immune escape capabilities. Being an essential mediator of infections and a key target of antibodies, mutations of its spike glycoprotein play a vital role in modulating its evolutionary trajectory. Here, we present a time-resolved statistical method, Dynamic Expedition of Leading Mutations (deLemus), to analyze the evolutionary dynamics of the SARS-CoV-2 spike. Together with analysis of its single amino acid polymorphism (SAP), we propose the use of ***L***-index in quantifying the mutation strength of each amino acid site, such that the evolutionary mutation pattern of the spike glycoprotein can be unravelled.

## I. Introduction

Ever since the emergence of coronavirus disease 2019 (COVID-19) in December 2019,^[1]^ its rampant propagation has greatly hampered global socioeconomic activities, all the while leaving billions infected and millions dead.^[2]^ Tremendous efforts have been put into mitigating the effects of this devastating pandemic by the international community, like implementing social lockdowns and mass vaccination programs. Regrettably, these measures have been ineffective in curtailing the persistence of this disease. The constantly shifting epidemiology of COVID-19 since its initial outbreak has been the result of the continuous evolution of its etiological agent, severe acute respiratory syndrome coronavirus 2 (SARS-CoV-2). Despite possessing one of the largest genomes among RNA viruses,^[3]^ coronaviruses exhibit comparatively lower substitution rates than their viral counterparts, as they are capable of encoding a 3′-to-5′ exoribonuclease located at their nonstructural protein 14 for RNA-proofreading,^[4–6]^ thereby facilitating proper genome replication by interacting with the low-fidelity RNA-dependent RNA polymerase.^[7,8]^ However, this proofreading mechanism is not flawless, as the exoribonuclease lacks the ability to repair nucleotide deletions, which translationally alters the amino acid sequences of viral proteins.^[9]^ Aside from the accumulation of deletions, another driving force by which coronaviruses evolve is RNA recombination.^[10,11]^ By admixing the genomic content of genotypically dissimilar coronaviruses, it is possible for novel phenotypes to be introduced into the viral recombinants.^[10,12]^ These evolutionary mechanisms have therefore enabled SARS-CoV-2 to amass a wide range of mutations as the pandemic progresses.

The immense selection pressures exerted upon SARS-CoV-2 have prompted its rapid genetic diversification, from which over 2500 lineages have been generated.^[13]^ Within the first two years of the COVID-19 pandemic, the World Health Organization (WHO) had already designated five SARS-CoV-2 variants as variants of concern, namely alpha (*α*, B.1.1.7), beta (*β*, B.1.351), gamma (*γ*, P.1), delta (*δ*, B.1.617.2), and omicron (*o*, B.1.1.529), as well as numerous other variants of interest.^[14]^ In rapid succession, each of these reported variants (RVs) emerged, proliferated, and outcompeted its antecedent, resulting in wavelike resurgences of COVID-19 cases. The emergence of all these variants has introduced many novel mutations that continue to enhance the fitness of the virus.^[15]^ Mutations of this nature, while uncommon,^[16]^ have substantially complicated the development of therapeutics, which greatly hinders the progress of COVID-19 treatment research.^[17]^ Hence, in order to develop more effective treatments and disease control strategies, it is of paramount importance to understand how this virus evolves over time.

Across the mutational landscape of the vast SARS-CoV-2 genome, the spike glycoprotein gene sits atop its plateau. The spike glycoprotein is a trimeric type I viral fusion protein that binds SARS-CoV-2 virion particle to the angiotensin converting enzyme 2 (ACE2) receptor on a host cell.^[18]^ In terms of its structure, the spike glycoprotein is composed of 2 subunits, the N-terminal subunit 1 (S1) and C-terminal subunit 2 (S2), within which multiple domains lie. The S1 region mainly facilitates ACE2 binding,^[19,20]^ and is made up of an N-terminal domain (NTD), a receptor-binding domain (RBD), and 2 C-terminal subdomains (SD1 and SD2), while the downstream S2 region is responsible for mediating virus-host cell membrane fusion.^[21]^ Being a key mediator of cellular infections via both cell surface and endosomal entry pathways,^[18]^ the spike glycoprotein is a primary target for antibodies,^[22,23]^ immune effector cells,^[24,25]^ and COVID-19 therapeutics.^[26–29]^ Due to these selection pressures, the spike glycoprotein exhibits much higher mutational activities than other SARS-CoV-2 structural proteins, whose mutations are capable of altering its infectivity and antigenicity.^[17,30]^ With its functional significance, elucidating the evolutionary trajectory of the spike glycoprotein would be the first step in deciphering the mechanisms behind virus evolution as a whole.

Nonetheless, even with extensive international collaborations in clinical and laboratory investigations, research on understanding SARS-CoV-2 spike glycoprotein evolution is still in its infancy. The complete sequence space of the spike glycoprotein, formed by all possible amino acid combinations of 1273 residues, encompasses more than 20^1000^ different sequences. However, its most recent evolutionary pathway envelops only a very small fraction of the entire sequence space, comprising a few million (*∼*10^6^) distinct sequences only. While data retrieved from Global Initiative on Sharing Avian Influenza Data (GISAID) reveals that nearly all spike glycoprotein residues have mutated at least once since the initial COVID-19 outbreak, only few of these amino acid sites exhibit polymorphism. In other words, most identified spike glycoprotein sequences are concentrated within a small region of the whole sequence space, and that majority of the possible sequences remain unexplored. This shows that, even though the evolutionary process of the spike glycoprotein is highly dynamic, particular mutational constraints exist to prevent the spike glycoprotein from navigating through all the possible sequence options. In this work, a method named Dynamic Expedition of Leading Mutations (deLemus) is developed to quantitatively characterize the robust properties of the SARS-CoV-2 spike glycoprotein evolutionary dynamics, from which variations in both mutation rate and single amino acid polymorphism (SAP) can be identified. With each domain of the spike glycoprotein possessing divergent functions that are crucial in facilitating the infection process of SARS-CoV-2,^[27]^ ***L***-index from our proposed deLemus analysis has been employed to quantify the mutational activity of each amino acid site, where it has been demonstrated that ***L***-index is capable of effectively outlining novel leading mutations (LMs) before their corresponding emerging variants are designated as RVs by the WHO.

## II. Methodology

Detecting dynamic patterns from big data sets has always been a major challenge in data analysis. In this work, we propose one method, dubbed as deLemus, to investigate the evolutionary dynamics of the SARS-CoV-2 spike glycoprotein at an amino acid sequence level.

A collection of 7,940,305 SARS-CoV-2 spike glycoprotein amino acid sequences was downloaded from the GISAID hCoV-19 database on February 9^th^ 2022.^[31]^ We used EPI ISL 402124 as the reference sequence of the spike glycoprotein.^[1,32]^ As there exists a substantial amount of repeated sequences in the original data, all degenerate sequences have been removed to keep only the non-degenerate ones for further analysis of sequence mutations. Overall, 172,280 non-degenerate sequences have been retrieved from the total set of reported sequences uploaded to GISAID between December 2019 and December 2021.

Sequences submitted within the same month were grouped together. Multiple sequence alignment was then consecutively conducted on each group using Clustal Omega to check the occurrence of substitution or deletion at each amino acid site,^[33]^ with respect to the reference sequence. This yielded the total number of mutated sites in all sequences ***n*** and the number of sequences *P* (***n***) with a given ***n*** in each month. The mutation rate Ξ in the unit of seq^-1^mo^-1^ was calculated based on the total number of mutations per sequence per month (Fig. S2). The total number of SAP at each *j*^th^ amino acid site in each *t*^th^ month ***s***_*j*_(*t*) and the number of amino acid sites *N* (***s***) were also computed, from which Poisson distribution was observed, giving the monthly average SAP number 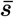 (Fig. S3).

For each *t*^th^ month, one *m ×n* mutation matrix ***H***(*t*) is constructed based on the multiple sequence alignment data, where *m* is the number of non-degenerate sequences displayed in a particular month, and *n* is the length of the spike glycoprotein amino acid sequence. In other words, each row represents one non-degenerate sequence from that month, and each column corresponds to one residue in the sequence. For the *i*^th^ sequence, if the *j*^th^ residue is changed, the corresponding matrix component ***H***_*ij*_(*t*) would be set to 1. Otherwise, it would be set to 0.

We then factorized ***H***(*t*) by singular value decomposition,^[34]^

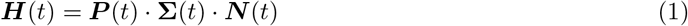

where ***P*** (*t*) is an *m* × *m* matrix and ***N*** (*t*) is an *n* × *n* matrix. (see Supplemental Material for details). From the monthly **Σ**_*i*_(*t*) and ***N***_*ij*_(*t*), we collected the top 4 leading sets of mutations to calculate the ***L***-index ***L***_*j*_(*t*), which is used to quantify the mutation strength of each *j*^th^ site in each *t*^th^ month,

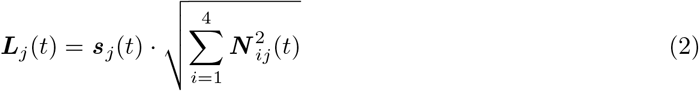

The amino acid sites were arranged according to their ***L***_*j*_(*t*), and top-ranked ones were identified as LMs of each month from January 2020 to December 2021.

## III. Results and Discussions

The structural information of each confirmed LM site was presented via AlphaFold2,^[35]^ as shown in Fig. 1. Here, we highlighted the mutation sites that have appeared in RVs. Based on the different characteristics of each protein domain, we divided the outlined LMs into four regions for discussion: NTD, RBD, SD1 and SD2, and S2. With the structural information provided in Fig. 1, functional changes of the spike glycoprotein conferred by these LMs can be illustrated.

**Fig. 1.**
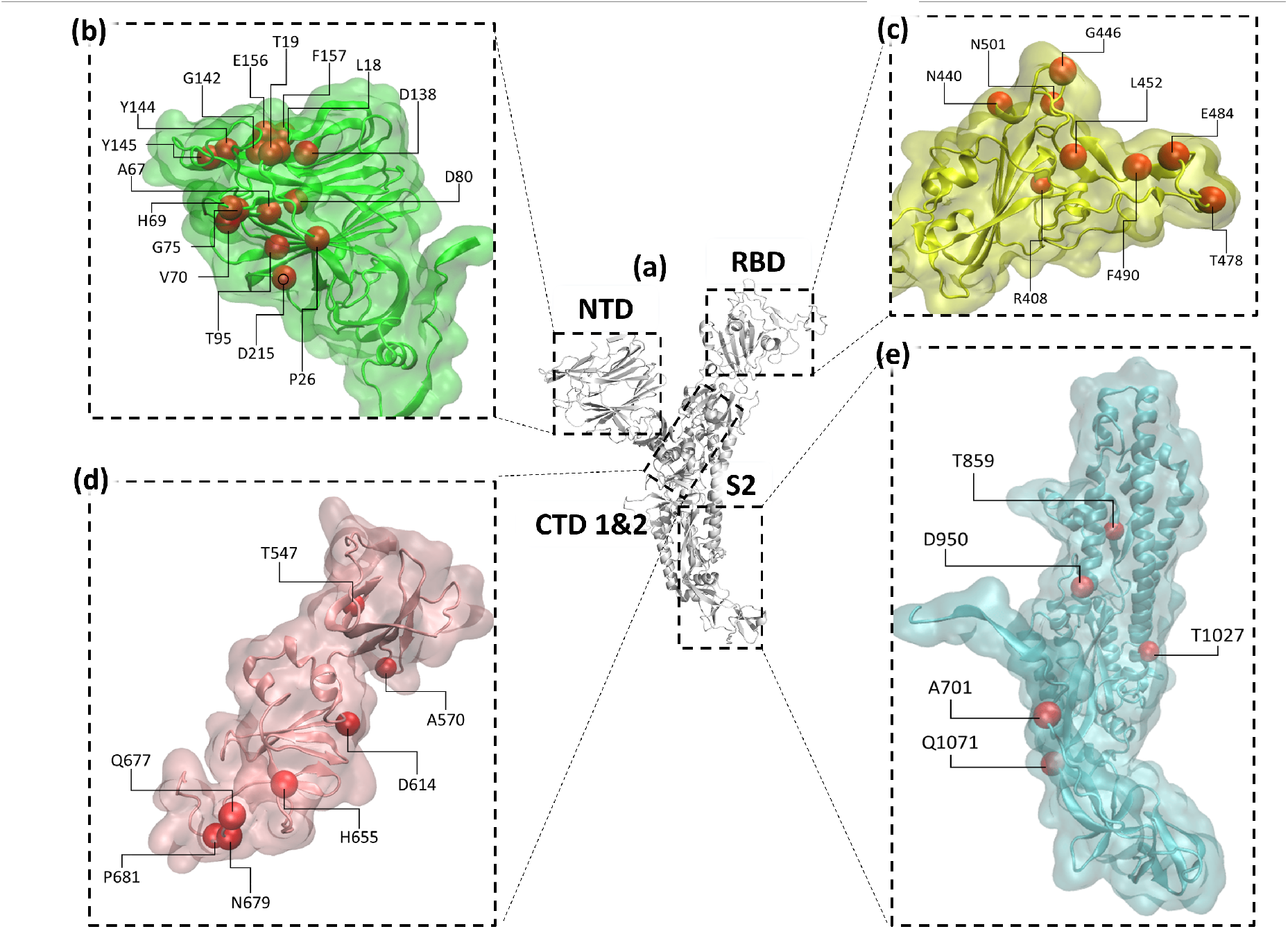
Confirmed LM sites (red spheres) in different domains of the SARS-CoV-2 spike glycoprotein. (a) A SARS-CoV-2 spike glycoprotein monomer (generated by AlphaFold2, Colab version, using the reference sequence EPI ISL 402124) encompassing the four major functional domains: NTD, RBD, SD1 and SD2, and S2. (b) Mutation sites outlined in NTD (green), most of which are clustered in the outer regions. (c) Mutation sites outlined in RBD (yellow), nearly all the captured mutations are located in the receptor-binding motif (RBM, 438-506). (d) Mutation sites outlined in SD1 and SD2 (pink). (e) Mutation sites outlined in S2 (cyan).

### A. N-Terminal Domain (NTD)

The NTD is an S1 ectodomain situated at the outermost region of the SARS-CoV-2 spike glycoprotein, on which several epitopes lie.^[16,36–38]^ While the NTD does not directly interact with ACE2 receptors, the domain’s close spatial proximity to the RBD has enabled some of its mutations to alter the cell entry dynamics of SARS-CoV-2.^[39,40]^ With their abilities to vastly impact the antigenicity and infectivity of the virus, it is of utmost importance to closely monitor the evolutionary trajectory of the NTD (Fig. 2).

**Fig. 2.**
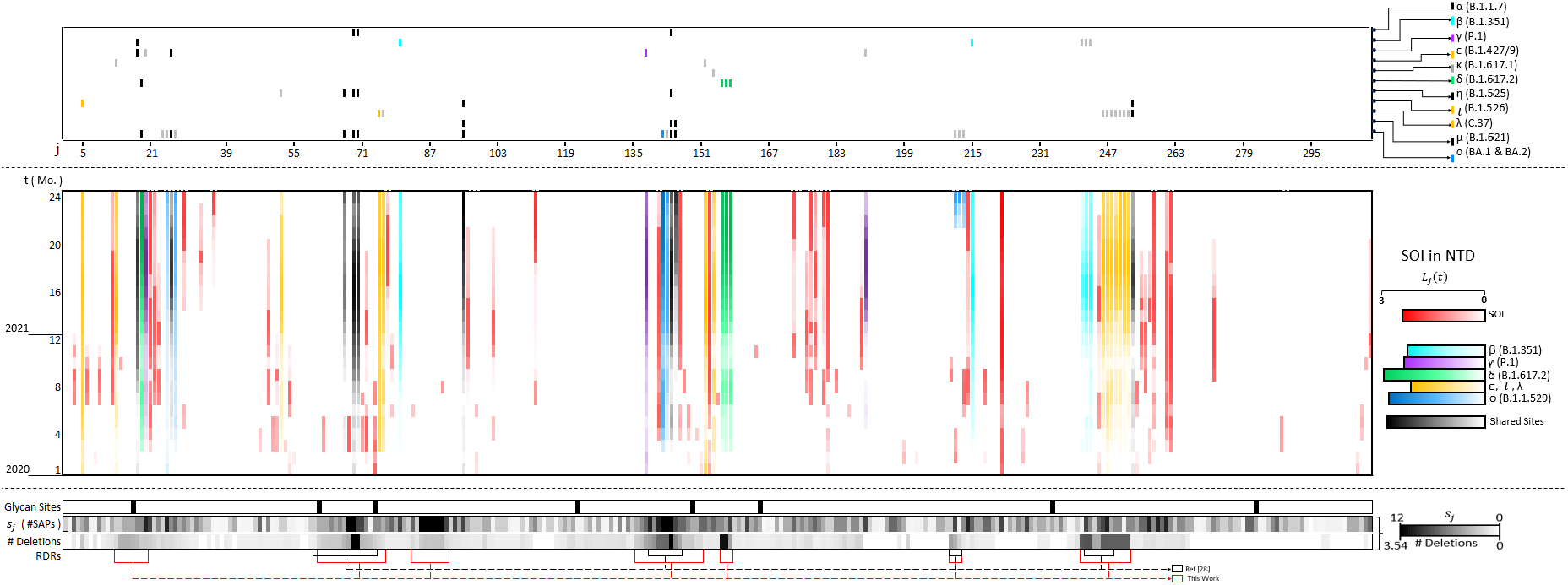
Evolutionary trajectory of the NTD between January 2020 (1^st^ mo.) and December 2021 (24^th^ mo.). Top panel: NTD mutations harbored by RVs as defined by the WHO. Sites shared by more than 1 RV are colored in black. Middle panel: NTD LMs outlined by our deLemus analysis, whose mutational activities are measured by their ***L***_*j*_(*t*). Bottom panel: Distributions of glycosylation sites, ***s***_*j*_(*t*), and deletions in the NTD, as computed by their mean values of occurrence in all non-degenerate sequences collected between March to August 2021. Bracketed stretches indicate the locations of recurrent deletion regions (RDRs), where those reported in ref^[9]^ are in black, and those identified in this work are in red.

Domains across the SARS-CoV-2 spike glycoprotein generally adopts substitution as a common mode for generating mutations, which would be discussed thoroughly in later parts. However, particularly in the NTD, insertions and deletions are also prevalent mechanisms by which mutations are generated, leading to its structural plasticity.^[16,41]^ Deletions in this domain have been characterized to frequently occupy particular locations of the spike gene known as RDRs, where partial nucleotide removals within specific stretches of codons can give rise to new nucleotide arrangements.^[9,42]^ One such example would be a persistent six-nucleotide deletion ΔAGTTCA within the spike gene segment GAGTTCAGA that results in the formation of the mutated codon CGA, replacing _156_EFR_158_ with a single glycine residue (R158G) in the *δ* variant. Interestingly, with regards to NTD deletions, data from our deLemus analysis shows that lengths of these RDRs expand over time (Fig. 2 and Fig. S5).

The overwhelming mutational activity of this domain can be clearly demonstrated by how the emergence of each RV introduces novel mutable sites in the NTD. Our investigation began with the ***L***-index calculation of each NTD amino acid site starting from December 2019 to outline potential LMs. By November 2020 (11^th^ mo.), we identified six LMs among numerous candidates, all of which were later confirmed to be present in the *α* and *β* variants that emerged in December 2020. These included all three mutations in the *α* NTD (ΔH69/V70 and ΔY144) and three mutations in the *β* NTD (L18F, D80A, and D215G), as presented in Fig. 1 and Fig. 2. The three *α* mutations are situated within previously documented RDRs (RDR1 and RDR2),^[9]^ where the former double deletion has been reported to promote infectivity by enhancing syncytia formation,^[43]^ while the latter single deletion located at the NTD N3 loop has been found to impair the neutralizing activities of several antibodies.^[44,45]^ As for the recorded *β* NTD mutations, all three of them have been found to confer immune escape capabilities.^[45,46]^

In December 2020 (12^th^ mo.), we identified multiple potential LMs in the NTD, two of which were later confirmed to be present in the *γ* variant that emerged in January 2021: P26S and D138Y. In the following month (13^th^ mo.), we outlined additional LMs, two of which, L5F and T95I, were then reported in the iota variant (*ι*, B.1.526) that emerged in February 2021 (14^th^ mo.). By the subsequent month (14^th^ mo.), we identified several other LMs, one of which was later confirmed to be present in the eta variant (*η*, B.1.527) that emerged in February 2021. While the functions of many of these mutations are not well-understood, it has been reported that the two LMs present in the *γ* variant disrupt the epitope of monoclonal antibody (mAb) 159, leading to a significant reduction in its neutralizing activity.^[47]^

Similarly, we identified multiple LMs in March 2021 (15^th^ mo.), three of which were later confirmed to be present in the *δ* variant that emerged in April 2021^[48]^: T19R and the ΔE156/F157 double deletion. Moreover, we identified a novel LM, G75V, which was later harbored by the lambda variant (*λ*, C.37) emerged in the same month as the *δ* variant. Interestingly, while the ΔE156/F157 double deletion has been experimentally shown to improve viral fitness by conferring immune evasion capabilities against NTD-targeting antibodies,^[49]^ the G75V mutation has been shown to significantly decrease the infectivity of SARS-CoV-2.^[50]^

We identified more novel LMs in October 2021 (22^nd^ mo.), two of which have been confirmed as novel NTD mutations carried by the first *o* sublineage, the BA.1 strain: ΔG142 and Y145D (Fig. 1 and Fig. 2). This variant first originated from South Africa in late November 2021.^[51,52]^ While the effects regarding mutations at G142 are not well-studied, its location within the NTD antigenic supersite implicates that G142 mutations may alter spike-antibody interactions,^[45]^ as one study has demonstrated that the G142D mutation is capable of conferring significant resistance against NTD-targeting mAbs.^[53]^ As for Y145D, this mutation has been reported to reduce the neutralization potencies of convalescent sera.^[54]^

### B. Receptor-Binding Domain (RBD)

The RBD is a S1 domain that not only plays an essential role in ACE2 recognition,^[55,56]^ but also acts as a region of immunodominance targeted by around 90% of all plasma or serum neutralizing antibodies.^[16,57,58]^ Mutations in this domain therefore commonly possess the abilities to alter virus-ACE2 or virus-antibody binding affinities,^[59]^ which enable the generation of variants with higher transmissibility or immune escape capabilities.^[16,60]^ In fact, as shown in Fig. 1, most mutations in this domain are located in the RBM that serves as a spike-ACE2 binding interface. These mutations would therefore undoubtedly affect the infectivity of the virus. With such significant functional implications, it is necessary to track RBD mutations in a temporal manner.

Since the emergence of the first two RVs, *α* and *β*, different SARS-CoV-2 variants have continued to acquire mutations in the RBD, including the latest *o* family. Several of these RBD mutations have been successfully captured by our deLemus analysis. One such important mutation is N501Y, which was first outlined as an LM (9^th^ mo.) by our deLemus analysis, confirmed in the *α* variant, and subsequently retained in the *β, γ*, mu (*μ*, B.1.621.1), and *o* variants, as depicted in Fig. 3. Studies have shown that the N501Y mutation can enhance ACE2 binding affinity by introducing *π*-*π* interactions between N501Y of RBD and Y41 of ACE2.^[47]^ In addition, this mutation has been found to decrease the neutralizing activities of some mAbs.^[46,60]^

**Fig. 3.**
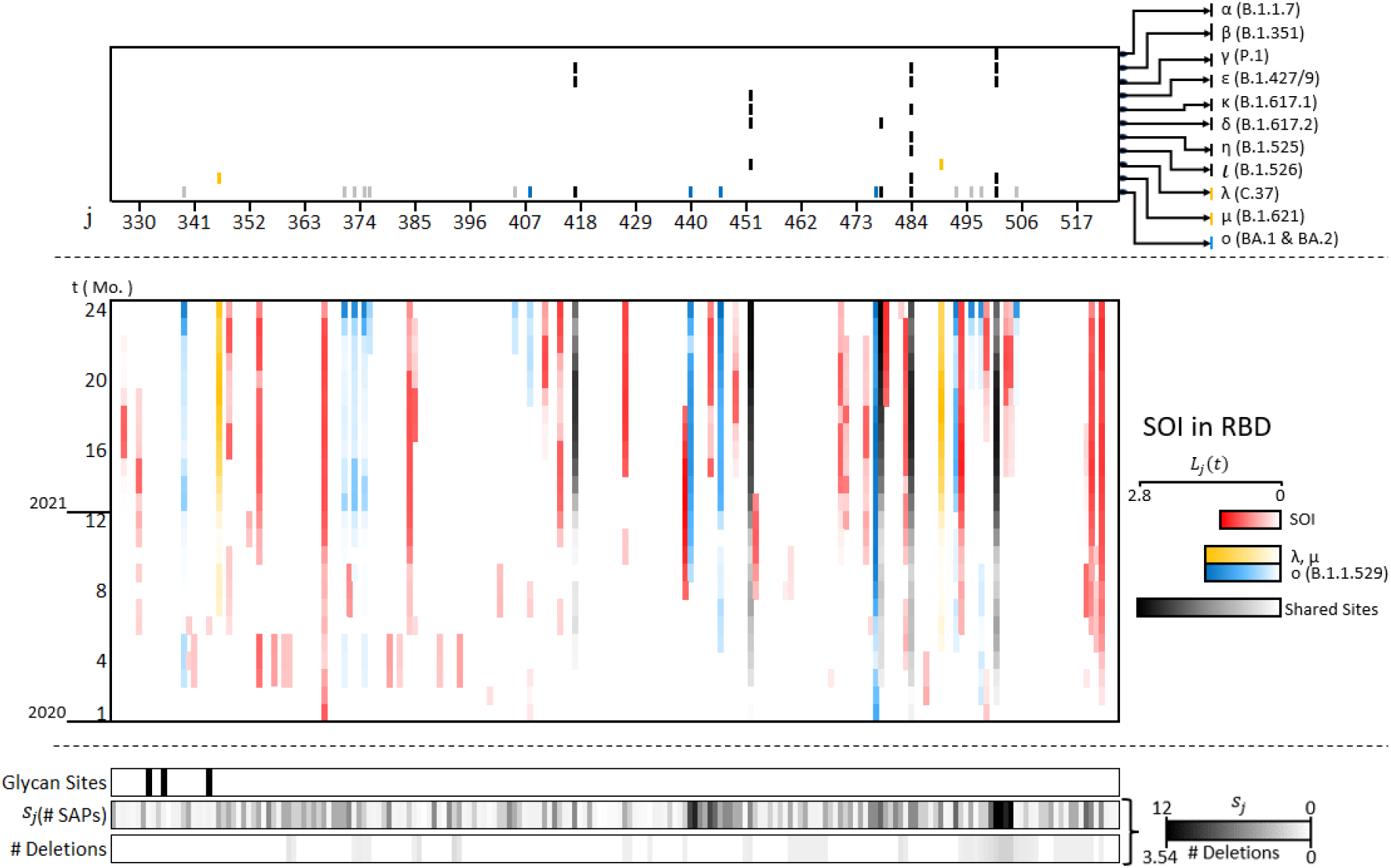
Evolutionary trajectory of the RBD between January 2020 (1^st^ mo.) and December 2021 (24^th^ mo.). Top panel: RBD mutations harbored by RVs as defined by the WHO. Sites shared by more than 1 RV are colored in black. Middle panel: RBD LMs outlined by our deLemus analysis, whose mutational activities are measured by their ***L***_*j*_(*t*). Bottom panel: Distributions of glycosylation sites, ***s***_*j*_(*t*), and deletions in the RBD, as computed by their mean values of occurrence in all non-degenerate sequences collected between March to August 2021.

E484K is another mutation that was first outlined as an LM (8^th^ mo.) by our deLemus analysis, and later confirmed in the *β* variant. Mutations in this site have been demonstrated to not only significantly reduce the neutralization titers of convalescent plasma, but also lower the activities of mAbs.^[61–63]^ Mapping of mutations has revealed that E484K is the strongest escape site for class 2 antibodies,^[64]^ which largely represent those found in convalescent polyclonal plasma. Although some studies suggested that mutations of the E484 residue would lead to diminished electrostatic complementarity between the RBM and the ACE2 receptor,^[65]^ many structural biology studies have shown that, when introduced with other mutations like the aforementioned N501Y, the E484K mutation can impart increased RBD-ACE2 binding. For instance, in the *β, γ* and *o* variants, the E484K-N501Y-D614G triple mutation is found to enhance RBD-ACE2 binding by inducing local rearrangements involving rotamer placements between Q493 of RBD and H34 of ACE2.^[66]^

Two other novel RBD mutations, L452R and T478K, were also outlined as LMs (11^th^ mo. and 13^th^ mo. respectively) by our deLemus analysis. These mutations are later confirmed as fingerprint mutations of the *δ* variant that emerged in April 2021. Their locations within the epitope of several important neutralizing antibodies enable them to enhance the immune escape capabilities of the virus.^[60,67–69]^ Furthermore, these two mutations have been reported to improve RBD-ACE2 interactions.^[70–72]^ For the former mutation, computational studies have shown that variants possessing the L452R-E484Q-N501Y triple mutation exhibit a secondary structure rearrangement that is associated with an increase in RBD-ACE2 binding affinity.^[73]^ As for the latter mutation, structural analysis has revealed that the T478K substitution allows the formation of two new hydrogen bonds located between Y489 of RBD and Y83 of ACE2, and F490 of RBD and K31 of ACE2 respectively, which results in tighter binding between the RBD and the ACE2 receptor.^[72]^

Known for its exceptionally high transmissibility, the *o* variant harbors a significant number of RBD mutations. Most of them have been reported in previous variants, but several new sites were outlined as LMs (22^nd^ mo.) by our deLemus analysis, which encompasses the mutations at R408, N440 and G446. The R408S mutation has been shown to alter the antigenic property of the spike glycoprotein by disrupting the binding of F2 antibodies.^[74]^ Unlike R408S, the N440K mutation can enhance spike-ACE2 binding affinity by increasing the electrostatic complementarity between the structurally flexible ACE2 recognition site and the ACE2 receptor.^[75]^ As for the G446 residue which is situated at a highly antigenic region of the spike structure (Fig. 1), changes at this site have been shown to influence neutralization by both mAbs and antibodies present in polyclonal serum.^[64,68,69]^

### C. Subdomains (SD1 & SD2) and S2 Subunit

The post-RBD region of the SARS-CoV-2 spike glycoprotein is consisted of SD1, SD2, and S2. While these regions do not directly engage the ACE2 receptor, they confer significant functions in spike allostery and post-ACE2-binding events respectively.^[76,77]^ For the two SDs, their close spatial proximities to the NTD-RBD linker motif enable them to modulate RBD motion, where mutations in these regions could alter RBD “up-down” equilibrium.^[78–80]^ S2, meanwhile, plays an important role in mediating host-virus membrane fusion by undergoing massive post-fusion conformational changes.^[77,81]^ Hence, mutations in S2 can drastically impact the fusogenicity of the SARS-CoV-2 spike, both in terms of host-virus membrane fusion for cell surface entry and cell-cell membrane fusion for syncytia formation.^[18,26]^ Additionally, the post-RBD region houses the S1/S2 and S2’ cleavage sites, which are proteolytically processed in order to facilitate membrane fusion processes.^[21,81–83]^ The former site in particular is unique to SARS-CoV-2 within the subgenus *sarbecovirus*, as it is generated from a polybasic insertion at the C-terminal of SD2 (_681_PRRAR ↓ S_686_), and has been experimentally found to be an important determinant of viral transmission.^[21,84]^ It is therefore quintessential to employ the deLemus method in tracking post-RBD mutations.

Like the previous sections, we first computed the ***L***-index of each post-RBD amino acid site in order to screen for potential LMs within SD1, SD2 and S2. By January 2020 (1^st^ mo.), we had already outlined D614G as one of the LMs, a prominent single amino acid substitution proximal to the SD1-SD2 junction which has emerged from as early as April 2020 (Fig. 1 and Fig. 4).^[85]^ Mechanistically, this mutation has been found to disrupt the interprotomer hydrogen bond involving S2 residues, thereby increasing the propensity for RBD to be in the “up” state.^[86]^ Experimentally, it has been shown that the D614G mutation enhances S1/S2 cleavage, cell entry, replicative fitness, and transmissibility.^[86–88]^ The increased viral fitness has therefore enabled D614G to be conserved in all RVs thereafter.

**Fig. 4.**
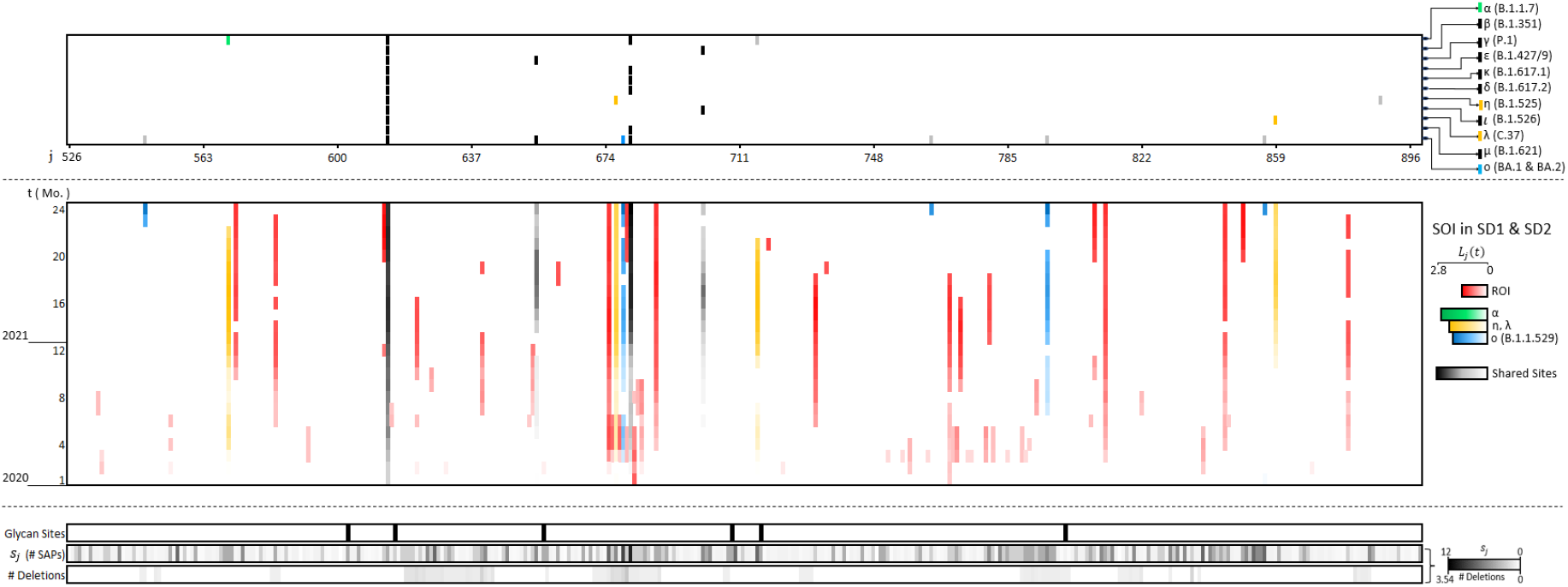
Evolutionary trajectory of the SDs between January 2020 (1^st^ mo.) and December 2021 (24^th^ mo.). Top panel: SD mutations harbored by RVs as defined by the WHO. Sites shared by more than 1 RV are colored in black. Middle panel: SD LMs outlined by our deLemus analysis, whose mutational activities are measured by their ***L***_*j*_(*t*). Bottom panel: Distributions of glycosylation sites, ***s***_*j*_(*t*), and deletions in the SDs, as computed by their mean values of occurrence in all non-degenerate sequences collected between March to August 2021.

In October 2020 (10^th^ mo.), we identified two LMs, A570D and P681H, both of which were later confirmed to be present in the *α* variant that emerged in December 2020. As shown in Fig. 1, these mutations are situated in the SD1 and SD2 region. In particular, the P681H mutation is located at the S1/S2 furin cleavage site and has been predicted to increase S1/S2 cleavability by substituting an electrically neutral proline with a cationic histidine, thereby enhancing the electrostatic complementarity between the cleavage motif and the anionic furin catalytic binding pocket.^[89,90]^ However, experimental studies have shown that this mutation only slightly increases local cleavage efficiency and does not significantly impact viral entry.^[91]^

In December 2020 (12^th^ mo.), we identified multiple LMs, two of which were later documented in two different RVs: the V1176F mutation was confirmed in the *γ* variant that emerged in January 2021; the Q677H mutation was confirmed in the *η* variant that emerged in February 2021. While the mutational effect of V1176F is not well-elucidated, the Q677H mutation of the *η* SD2 has been found to promote syncytia formation.^[92]^ In addition to these novel mutations, the amino acid site 681, which was initially characterized by its high ***L***-index in April 2020, developed substantial SAP by early 2021 (Fig. 4). This was demonstrated by its substitution to various basic residues. For instance, instead of retaining *α* variant’s P681H mutation, both the kappa (*κ*, B.1.617.1) and *δ* variants developed the novel P681R mutation. Unlike its predecessor, this mutation greatly improves furin cleavage efficiency and hence spike fusogenicity.^[93,94]^ We identified several more novel LMs within these domains in the following months, some of which were harbored by forthcoming RVs. These include the D950N mutation of the *δ* S2 and the T859N mutation of the *λ* S2 (Fig. 4 and Fig. 5).

**Fig. 5.**
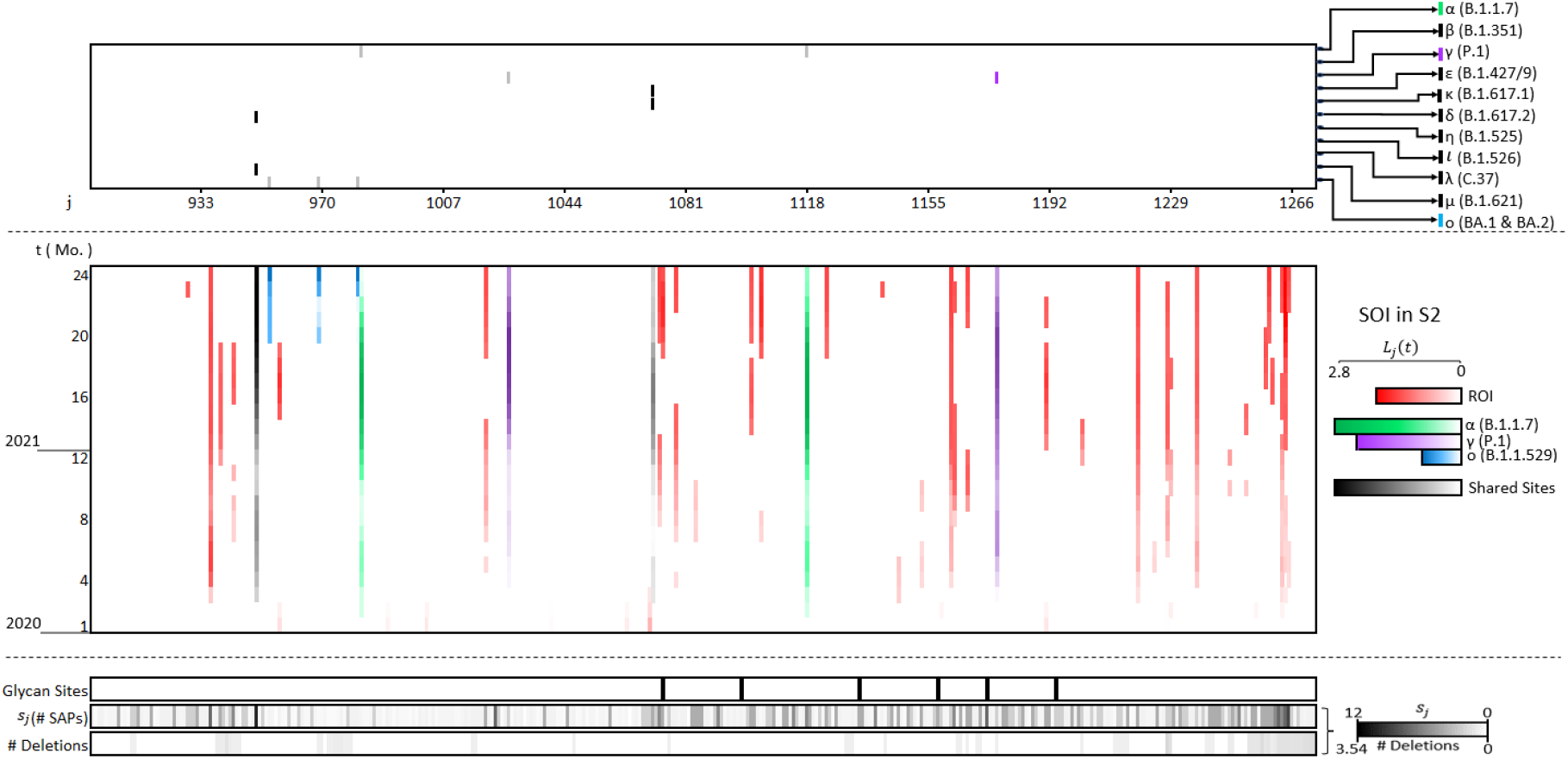
Evolutionary trajectory of the S2 between January 2020 (1^st^ mo.) and December 2021 (24^th^ mo.). Top panel: S2 mutations harbored by RVs as defined by the WHO. Sites shared by more than 1 RV are colored in black. Middle panel: S2 LMs outlined by our deLemus analysis, whose mutational activities are measured by their ***L***_*j*_(*t*). Bottom panel: Distributions of glycosylation sites, ***s***_*j*_(*t*), and deletions in the S2, as computed by their mean values of occurrence in all non-degenerate sequences collected between March to August 2021.

In October 2021 (22^nd^ mo.), we identified additional novel post-RBD LMs. One of these sites, N679K, was later confirmed to be present in the SD2 of the *o* lineage, shared by both the BA.1 and BA.2 strains (Fig. 4). This mutation had previously been detected in January 2021 (13^th^ mo.). The N679K mutation introduces a basic lysine residue near the S1/S2 furin cleavage site. However, experimental studies have suggested that, like the P681H mutation of *α*, N679K does not appreciably improve proteolytic processing of the spike glycoprotein.^[95]^

Aside from the outlined LMs, we also revealed some general evolutionary features of the SARS-CoV-2 spike glycoprotein, which can be quantified by Ξ and 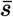 respectively. Mutations are the primary source of genetic variation; how mutation rates fluctuate over the course of evolution is of particular concern.^[96]^ In terms of its mutation rate, 3 characteristic phases of Ξ evolution can be distinguished throughout the pandemic. In the first phase between the initial outbreak of SARS-CoV-2 in December 2019 and October 2020, Ξ maintained a steady state at a relatively low value. A month before the emergences of *α* and *β* in December 2020, the second phase proceeded, when Ξ gradually increased until attaining a maximum value in March 2021, coinciding with the emergence of *δ*. Afterwards, the third phase followed, which marked the gradual decrease of Ξ up till June 2021, as it returned to a steady state comparable to that noted in the first phase. As most mutations on the spike glycoprotein are assumed to be deleterious, selection often acts against high mutation rates,^[97]^ which can be illustrated by the persistently low Ξ in the first evolutionary phase. Additionally, the stable Ξ in this phase may implicate the establishment of a dynamic equilibrium within a heterogeneous viral population,^[98,99]^ where each pre-*α* variant harbors similar levels of fitness. The subsequent rise in Ξ is thought to be a consequence of environmental changes that perturbed the equilibrium population of SARS-CoV-2. For better adaptability, elevating Ξ via the gain of a mutator allele would provide increased chances from which beneficial mutations would arise.^[100]^ However, more deleterious mutations would likewise be introduced if high Ξ value remains sustained.^[97,101]^ This would explain the gradual reduction of Ξ upon the start of the third phase. On the other hand, while genetic diversity measured by 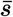 also exhibits slight fluctuations over the course of this pandemic, a clear increasing trend can be observed since December 2019. This growing 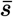 indicates the ongoing adaptive diversification of the SARS-CoV-2 spike glycoprotein.^[102]^ Nonetheless, further research would be necessary to elucidate the exact mechanisms that govern the complex evolutionary trajectory of SARS-CoV-2.

## IV. Conclusion

Our deLemus analysis allows the systematic monitoring of dynamic protein sequence samples, by extracting crucial information from time-dependent big data sets that accounts for the mutational activity of each amino acid site. Based on the variant decomposition data shown in Fig. S4, we defined the emergence of the *α, δ*, and *o* variants as the three most prominent phases of the COVID-19 pandemic. LMs outlined by our deLemus analysis were therefore compared against the reported mutations of these three RVs, as summarized in Fig. S8. Prior to the outbreak of the *α* variant, we captured seven of its ten reported mutations. Moving on to the *δ* variant, we were able to outline eight of its nine mutational sites as LMs. Lastly, we confirmed half of the mutational sites carried by the *o* variant. Overall, around 70% of the aforementioned RV mutations correspond to our outlined LMs, meaning that our deLemus analysis is capable of effectively capturing most RV mutations as LMs from the enormous data sets. Hence, these outlined LMs can serve as a guideline for deciphering the complex mutational pathways taken by the SARS-CoV-2 spike glycoprotein. Moreover, our deLemus analysis enables us to comprehend the time-resolved mutation patterns in each functional domain of the spike glycoprotein. For the NTD, our results verified the presence of previously characterized RDRs,^[9]^ and showed that these RDRs evolve by expansion over time, as indicated by their monthly ***L***-index computations depicted in Fig. 2. This highlights the importance of deletions in the evolutionary process of SARS-CoV-2. As well, our results showed a high occurrence of SAP in the NTD, particularly in common mAb-targeted regions.^[45,103]^ Collectively, the disproportionately high mutation rate of the NTD implies that the virus favors evolutionary routes in which both optimal ACE2-binding affinity and domain flexibility for immune escape are retained. For the RBD, we observed that most mutations harbored by previous RVs were incorporated into the RBD of the *o* variant, likely because these mutations are linked to fitness-enhancing traits. Given the RBD’s crucial role in mediating viral infections, which already shows robust binding affinity toward ACE2 receptors, we propose that the emergence of LMs in this region is driven by immune selection pressure. As a result, RBD mutations are likely to enhance viral transmissibility and antibody resistance.^[53]^ The rapid evolution of the SARS-CoV-2 spike glycoprotein is evident from its increasing number of mutated amino acid sites ***n*** and monthly SAPs 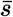 over time, as suggested by our deLemus analysis. To develop better pandemic control measures, it is quintessential to first understand how SARS-CoV-2 evolves. Therefore, sustained global efforts have to be called for a more comprehensive virus surveillance system.

## Supporting information

Supplemental Results

## V. Acknowledgements

We are grateful to Professor Yong Huang for many thought-provoking discussions and encouragement in the course of this work, which is supported in part by Hong Kong Branch of South Marine Science and Engineering Guangdong Laboratory (SMSEGL20SC01) (for H.S.), HKUST grant (R9418), ASPIRE League Partnership Seed Fund Program (ASPIRE 20191), and Society of Interdisciplinary Research (SOIR É E). This research made use of the computing resources of the X-GPU cluster supported by the Hong Kong Research Grant Council Collaborative Research Fund (C6021-19EF). We would also like to acknowledge the enlightening advice from Jie Gao in the early stage of this work; the technical support from Long T. Ta and Siqi Lei for improving figures; Ruichen Li, Leon R. Salim, and Salina Lam in data collection and preparation.

