## Supplemental Results for "Dynamic Expedition of Leading Mutations in SARS-CoV-2 Spike Glycoproteins"

(Dated: April 7, 2023)

### Contents

**Methodology of Dynamic Expedition of Leading Mutations (deLemus)** 2

**Time evolution of leading mutations** 3

**Time dependent distributions of $P(n)$ & $N(s)$** 5

**Time-dependent and cumulative distributions of $N(s)$** 6

**Variant decomposition of non-degenerate sequences** 7

**The average mutated sites per sequence over time** 8

**Evolution of deletion regions in the NTD** 9

**Distribution of deletion occurrences and $s$ (number of SAPs) in the NTD** 10

**RDRs and related mutations in VoC/VoI** 11

**Confirmed LMs** 12

**Comparison of LMs with VoCs** 13

---

[*] These authors contribute equally.

### Methodology of Dynamic Expedition of Leading Mutations (deLemus)

With the multiple sequence alignment data we collected, we constructed one mutation matrix $\boldsymbol{H}(t)$ for each month from January 2020 to December 2021, where $t$ represents the $t^{th}$ month starting from January 2020. For a month that has $m$ new individual sequence, the system is represented by an $m \times n$ matrix $\boldsymbol{H}(t)$. Different rows of the matrix represent sites mutation information in each sequence. Then, the sequence correlation matrix writes:

$$\boldsymbol{C}_{seq}(t) = \boldsymbol{H}^T(t) \cdot \boldsymbol{H}(t) \tag{1}$$

where $\boldsymbol{C}_{seq}(t)$ is an $n \times n$ matrix with $n$ eigenvectors. The matrix composed by eigenvectors of the sequence correlation matrix writes:

$$\boldsymbol{N}^T(t) = [\boldsymbol{N}_1^T(t), \boldsymbol{N}_2^T(t), \ldots, \boldsymbol{N}_n^T(t)] \tag{2}$$

Moreover, $\boldsymbol{H}(t)$ can be factorized according to SVD[1] as:

$$\boldsymbol{H}(t) = \boldsymbol{P}(t) \cdot \boldsymbol{\Sigma}(t) \cdot \boldsymbol{N}(t) \tag{3}$$

where $\boldsymbol{P}(t)$ and $\boldsymbol{N}(t)$ share the same eigenvalue series $\boldsymbol{\Sigma}(t)$. The contribution of the $i^{th}$ eigenvector of $\boldsymbol{N}(t)$ to the system is $\sigma_i^2(t)/\sum_i \sigma_i^2(t)$. Then, we defined the site mutation series $\boldsymbol{N}_{ij}(t)$ for each month as:

$$\boldsymbol{N}_{ij}(t) = (\boldsymbol{\Sigma}(t) \cdot \boldsymbol{N}(t))_{ij} \tag{4}$$

$\boldsymbol{N}_{ij}(t)$ represents the mutation amplitude of $j^{th}$ site corresponding to the $i^{th}$ eigenvector. Applying SVD on these matrices, we calculated the eigenvalue series $\sigma_i$ and mutation series $\boldsymbol{N}_{ij}(t)$ of SARS-CoV-2 as shown below. Here, we only kept the top 4 $\boldsymbol{N}_{ij}$ series. For each mutation set $i$, we defined the $\boldsymbol{L}_{ij}$ index of each mutation set as the product of its number of SAPs and mutation amplitude:

$$\boldsymbol{L}_{ij}(t) = \boldsymbol{s}_j(t) \cdot \boldsymbol{N}_{ij}^2(t) \tag{5}$$

The time evolutions of the 4 leading $\boldsymbol{L}_{ij}$ index sets are shown in Fig. S1(a), where different variants are labeled by different colors. The contribution of $\boldsymbol{L}_{ij}(t)$ to the mutation index of SARS-CoV-2 is $\sigma_i^2(t)/\sum_i \sigma_i^2(t)$. To get the overall effect of each SARS-CoV-2 spike mutation, we defined the $\boldsymbol{L}$-index of each $j^{th}$ site as the magnitude of the summed index of the top 4 mutation sets. In other words, the $\boldsymbol{L}$-index of the $j^{th}$ site for the $t^{th}$ month is defined as:

$$\boldsymbol{L}_j(t) = \sqrt{\sum_i^4 \boldsymbol{L}_{ij}^2(t)} = \boldsymbol{s}_j(t) \cdot \sqrt{\sum_{i=1}^4 \boldsymbol{N}_{ij}^2(t)} \tag{6}$$

Dynamic expeditions of LMs are presented in Fig. S1(a). Composition of the top 6 LM sets is shown in Fig. S1(b), where we can see the contribution of $5^{\text{th}}$ and $6^{\text{th}}$ mutations sets are already negligible. Thus, only the top 4 mutation sets are considered in this work.

### Time evolution of leading mutations

(a)

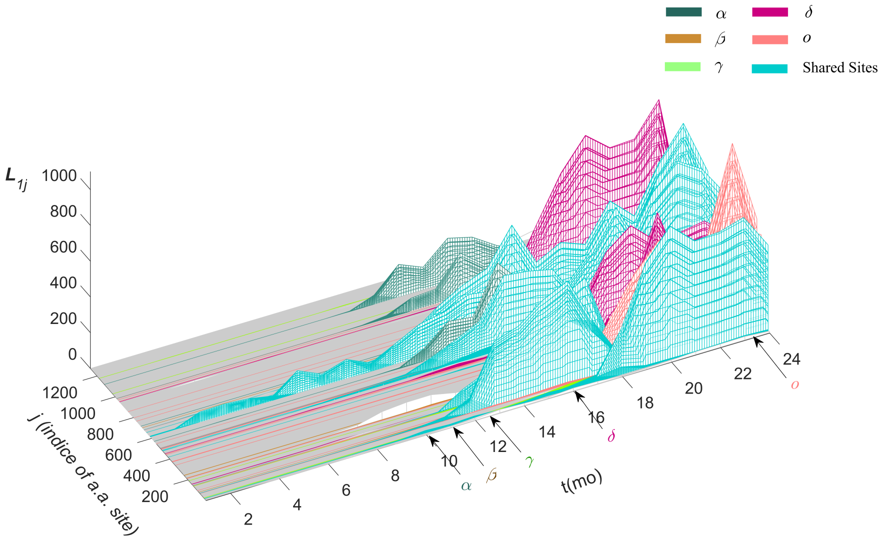


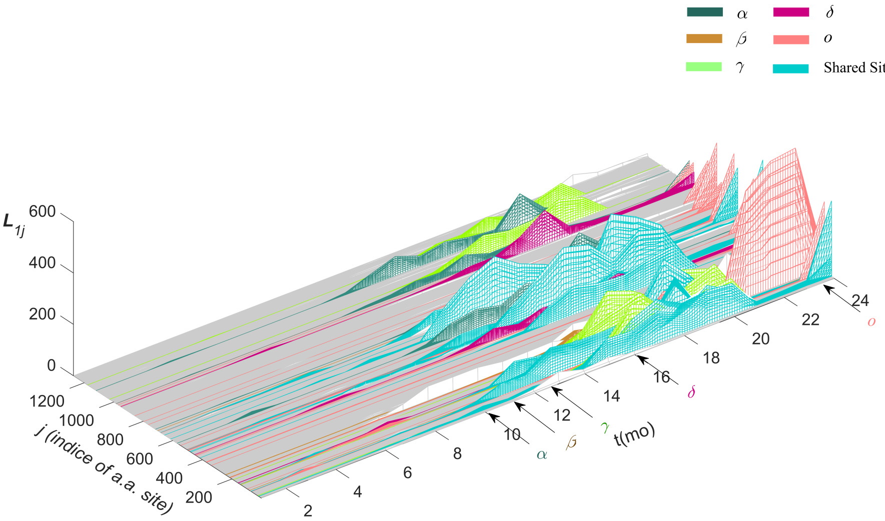


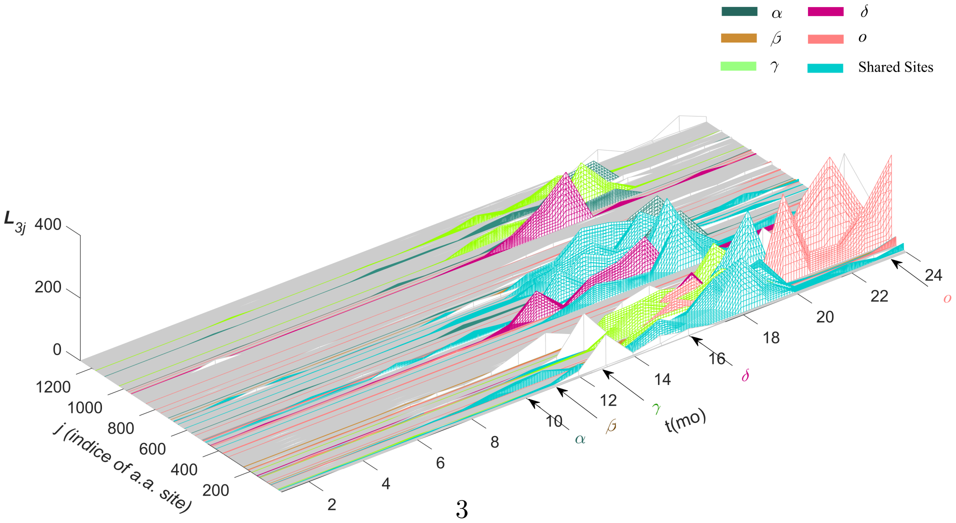

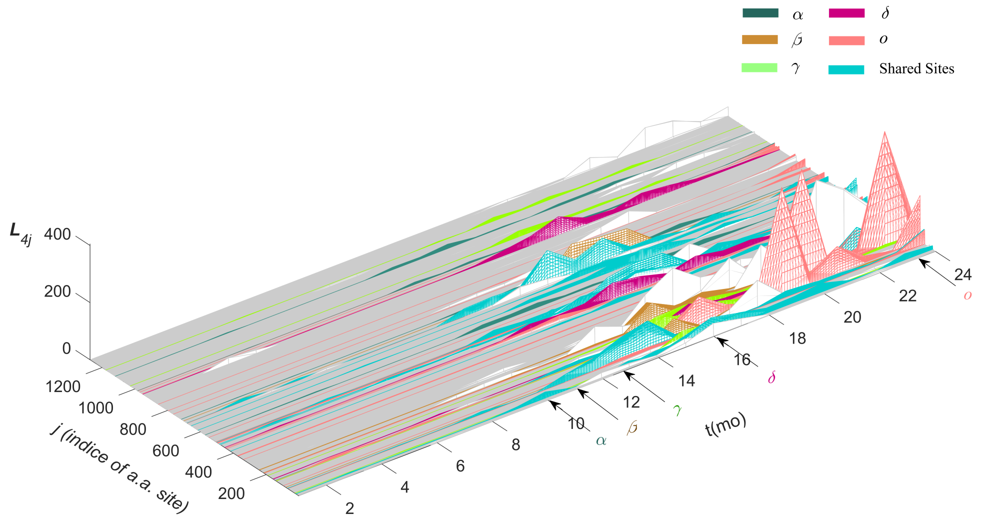


(b)

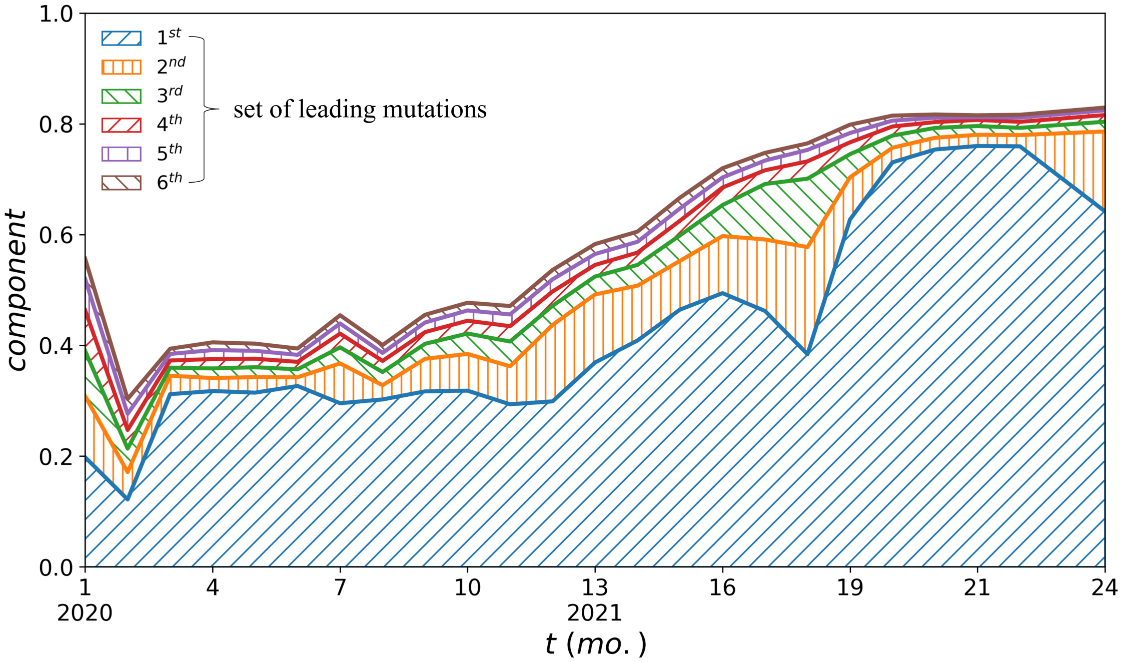


Fig. S1. (a). Time evolution top 4 mutation series. Shortly after the outbreak of COVID-19, amino acid site 614 quickly mutated and became the most common mutation from January to October 2020. In early October 2020, $\alpha$ variant-related sites started to become more mutationally active, which is indicative of the potential emergence of a new variant. After that, the $\alpha$ variant emerged, which quickly became the most widely spread SARS-CoV-2 lineage throughout the winter of 2020. Although the $\beta$ and $\gamma$ were reported in November and December 2020 respectively, mutational activities of sites specific to these two variants are suppressed by those in the dominant $\alpha$ variant. Hence, they did not show up until the second mutation set. At Apr. 2021, sites related to $\delta$ strain shows up and become the dominant strain in the first mutation set soon after it was reported, indicating the appearance of a new variant. Moreover, sites of VoC also show up but with lower mutation amplitude score. (b). Composition of top 6 leading mutation sets

### Time dependent distributions of $P(n)$ & $N(s)$

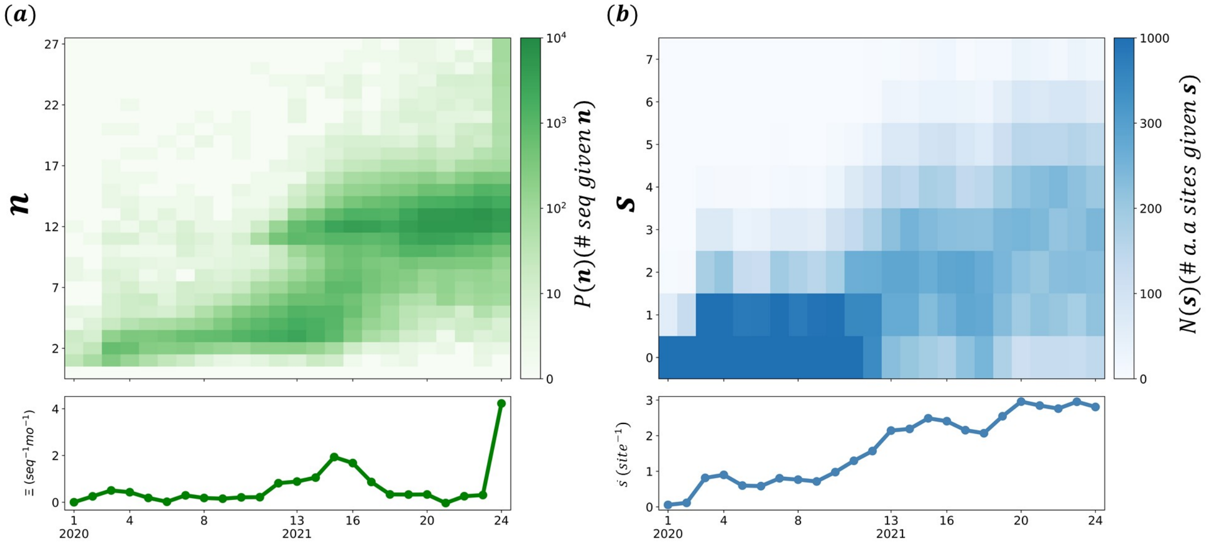


Fig. S2. (a) Time-resolved evolution of single-residue mutations in spike protein sequences. The top heat map denotes the time-dependent distribution of the number of mutations per sequence $n$. The bottom curve shows the time course of average mutation rate $\Xi$. (b) Time-resolved evolution of SAPs. The top heat map denotes the time-dependent distribution of the number of different amino acid substitutions at each site $s$. The bottom curve shows the time course of average substitution number $\bar{s}$.

#### Time-dependent and cumulative distributions of $N(s)$

(a)

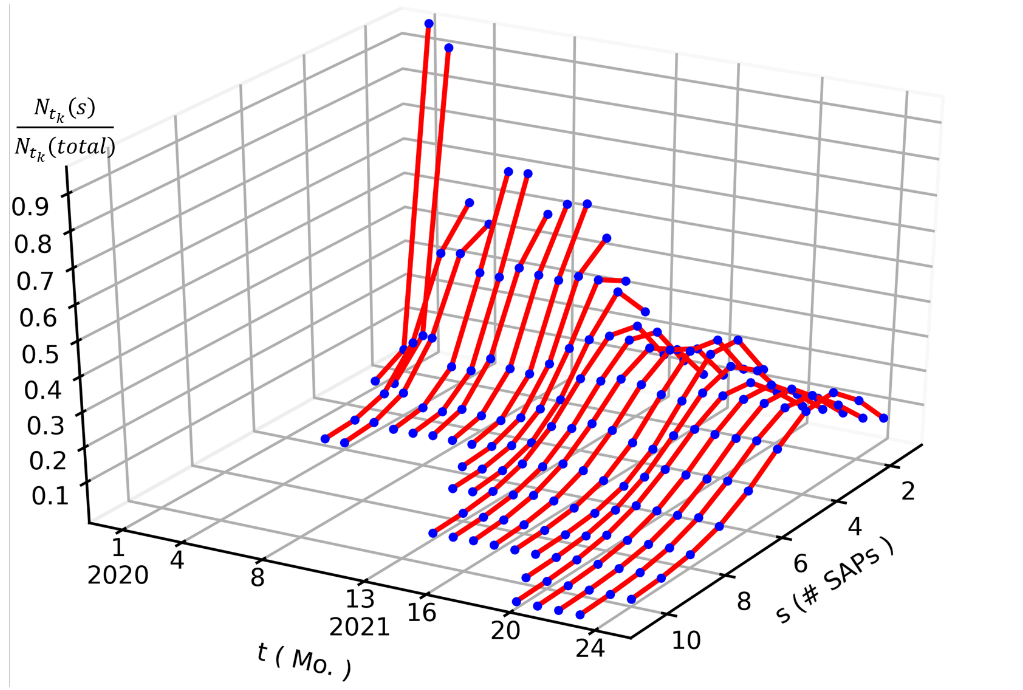


(b)

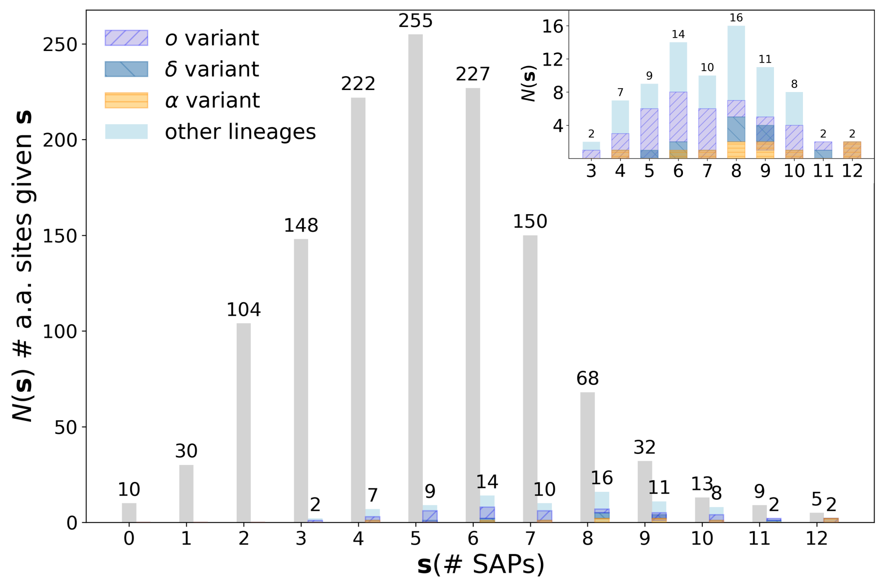


Fig. S3. (a). The dynamic of $N(\boldsymbol{s})$ distribution over the months. (b). Overall $N(\boldsymbol{s})$ distribution inside the time window from Jan. 2020 to Dec. 2021. The $N(s)$ is plotted in grey color, VoC/VoI is showed in blue. The composition of different strain is showed at top right corner of the plot with different color: purple for $o$, dark blue for $\delta$ and yellow for $\alpha$. Both $\alpha$, $\delta$ and $o$ show a preference of the region with more SAP options. Most VoC/VoI sites of SARS-CoV-2 are located on the right side of the mean value and nearly half of the sites have been reported as VoC/VoI exhibit a $\mathbf{s}$ value large than 9.

#### Variant decomposition of non-degenerate sequences

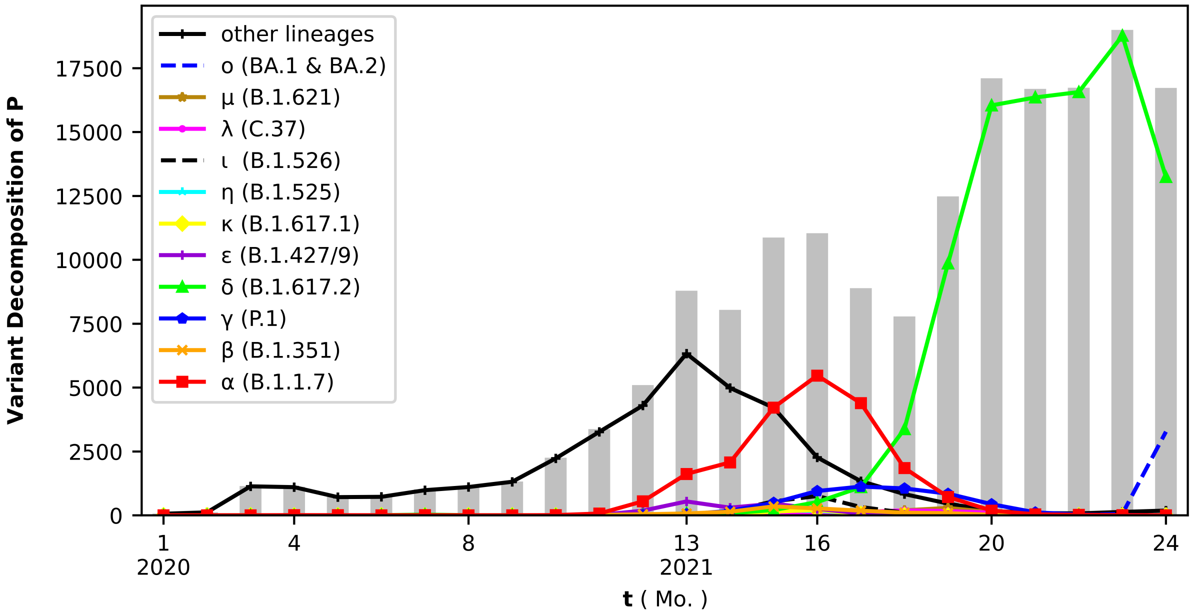


Fig. S4. Distribution of non-degenerate sequences are plotted in grey. Decomposition of all sequences is shown as colored lines. $\alpha$ and $\delta$ variants are the dominant lineages. The composition of lineages is indicative of the diversity of SARS-CoV-2 at different time points, similar to the trend observed in Fig. 1(b).

#### The average mutated sites per sequence over time

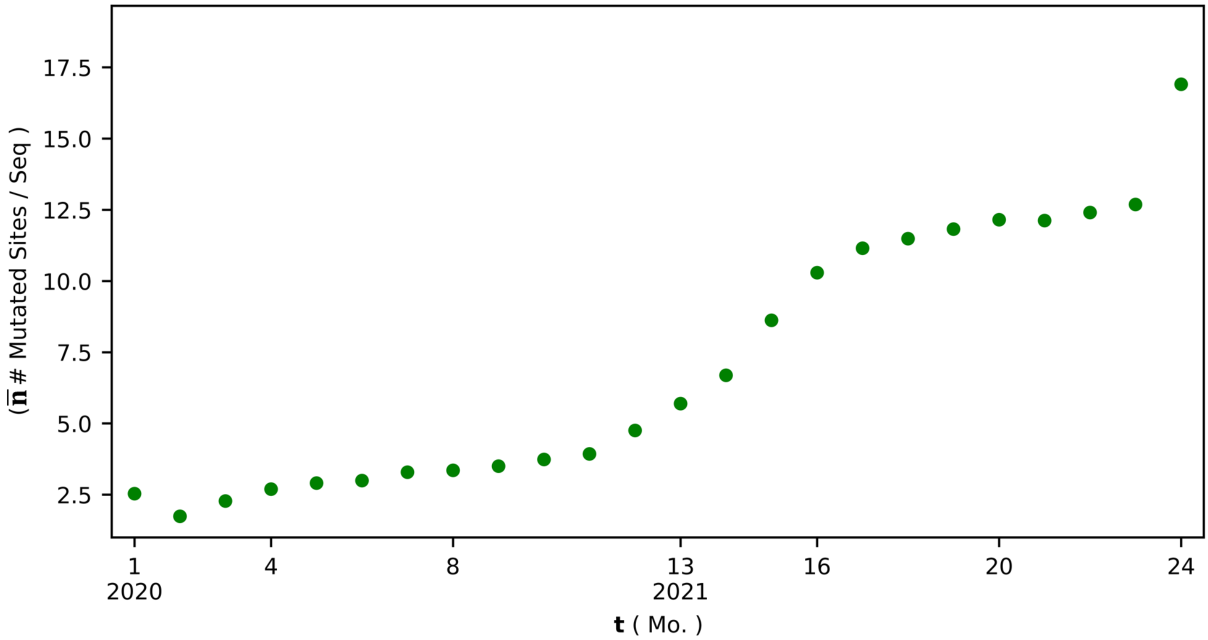


Fig. S5. $\bar{\boldsymbol{n}}_{t_k} = (\sum_{ij} \boldsymbol{H}_{ij})/m(t_k)$, where $\boldsymbol{H}_{i,j}(t_k)$ is the mutation matrix and $m(t_k)$ is number of sequences in the $k^{th}$ month. $\bar{\boldsymbol{n}}$ reflects the ensemble average deviation of all sequence compared to the initial sequence, which suggests the increments of mutated amino acid sites per month is around $0.5\tilde{2}$ (Fig. 1(a)). Additionally, the curve is observed to exhibit an accelerated growth between January and April 2021, which is indicative of the start of a new evolutionary stage for SARS-CoV-2, the dominance of $\alpha$ and $\delta$ variants.

#### Evolution of deletion regions in the NTD

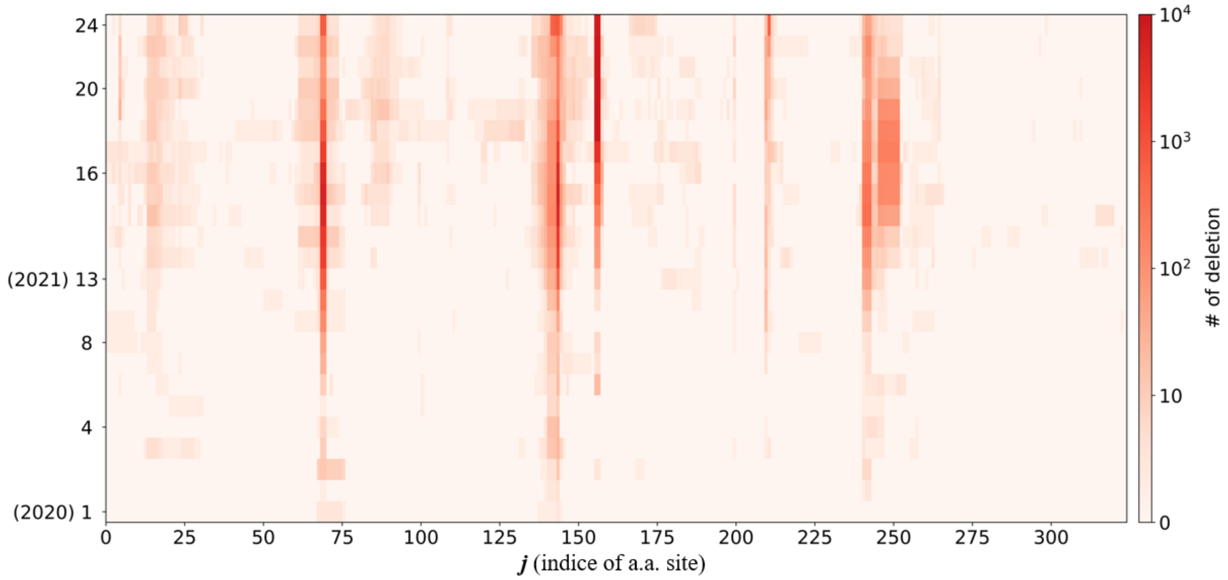


Fig. S6. Evolution of deletion regions in the NTD. Not only are all four of the previously identified RDRs ($\Delta 69-70$ (N2 loop), $\Delta 141-144$ (N3 loop), $\Delta 210$ (between N4 and N5 loop), and $\Delta 243-244$ (N5 loop)[3]) found to exhibit expansion over time, emergence of novel RDRs has also been observed, which encompass 3 more NTD RDRs, 1 RBD RDR, and 1 RDR proximal to the S1/S2 furin cleavage site (Table S1).

#### Distribution of deletion occurrences and $s$ (number of SAPs) in the NTD

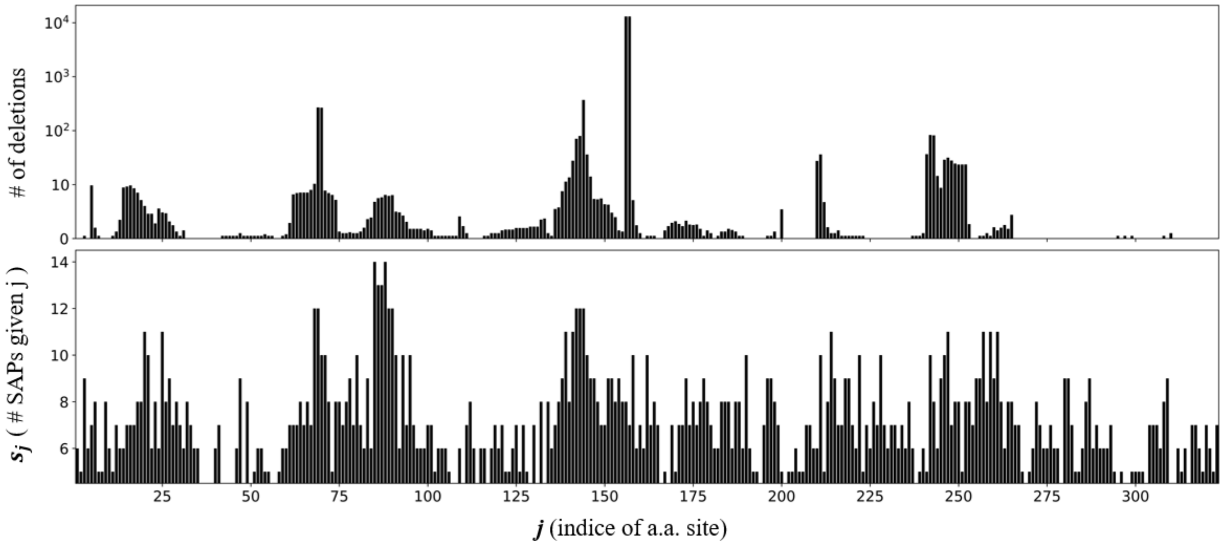


Fig. S7. Top panel: Distribution of deletion occurrences in NTD; Bottom panel: Distribution of $\mathbf{s}$ (Number of SAPs). As we can see, the peaks of both panels are located at similar positions. This indicates that the occurrences of deletions and SAPs exhibit positive correlation within the NTD, which suggests the presence of a common selection pressure that acts on those particular regions.

### RDRs and related mutations in VoC/VoI

| RDRs | Related Mutations in VOC/VOI |
|---|---|
| 13-27 | L18F(β,γ) T19R(δ) **T20N**(γ) P26S(γ) Δ24-26&A27S (BA.2) |
| 61-76 | A67V(η,o) Δ69-70(α,o,BA.2) G75V&T76I:λ |
| 83-91 | |
| 136-151 | D138Y(γ) Δ142-143&Y145D(o) Δ144(α,o,BA.2,η) Y144S&**Y145N**(μ) |
| 156-158 | Δ156-157 and R158G |
| 210-212 | Δ211&L212I(o) |
| 241-252 | Δ241-243(β) Δ246-252&**D253N**(λ) D253G(ι) |
| 500-504 | N501Y (α,β,γ,o,BA.2,μ) Y505H(o,BA.2) |
| 676-680 | N679K(o,BA.2) P681H(a,o,BA.2) P681R(δ,κ) |

| Shared epitopes |
|---|
| 14-20 |
| 140-158 |
| 245-264 |

| Glycan sites |
|---|
| N17, N61, N74, N122, N149, N165, N234, N282, N331, N334, N343 |

TABLE S1: Left panel: RDRs and related mutations in VoC/VoI. The first 4 RDRs match well with those characterized in previous work, with some extended regions[3]. The time evolution of RDRs can been seen in Fig. S6; Right panel: Top: Shared epitopes regions as reported in previous studies[4][5], which highly overlap with RDRs. Bottom: Glycan sites in the NTD. the shared epitopes regions sre broderred by four glycan sites N17, N74, N122 and N149.[5][6] T20N, Y145N and D253N are bold as these mutations may introduce glycosylations.

#### Confirmed LMs

| Confirmed SOIs | | |
|---|---|---|
| NTD | RBD | S2 |
| L5:ι | R346:μ | 614:All |
| L18:β,γ | R408:o | 677:η |
| T19:δ,o | N417:β,γ,o | 679:o |
| P26:γ,o | N440:o | 681:α,δ,κ,μ,o |
| A67:η,o | G446:o | 716:α |
| H69:α,η,o | L452:δ,κ,ε,λ | 859:λ |
| V70:α,η,o | S477:o | 950:δ,μ |
| G75:λ | T478:δ,o | 1176:γ |
| D80:β | E484:β,γ,κ,η,ι,μ,o | |
| T95:ι,μ,o | F490:λ | |
| D138:γ | N501:α,β,γ,μ,o | |
| G142:o | | |
| Y144:α,η,μ,o | | |
| E156:δ | | |
| F157:δ | | |
| D215:β | | |

TABLE S2: Confirmed LMs. Variants possessing these mutational sites are labeled by their corresponding Greek letters.

#### Comparison of LMs with VoCs

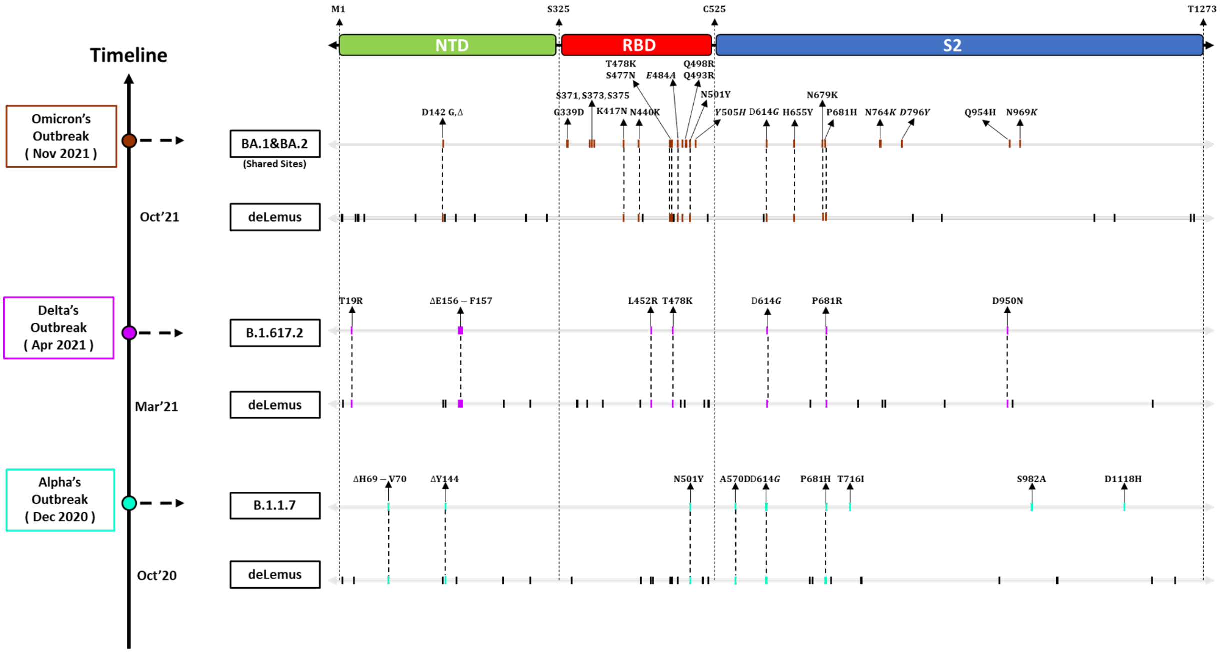


Fig. S8. Comparison of LMs with Variants of Concern. We tried to capture the spike mutations in variants of concern by using only the GISAID amino acid sequence data uploaded prior to their corresponding outbreaks. Based on the data collected until October 2020, we were able to capture 70% of the mutational sites carried by the $\alpha$ variant that emerged in December 2020, as shown in the above figure. By reapplying the same scheme to the $\delta$ and $o$ variants, it is revealed that our deLemus analysis has the prediction accuracy of 88% and 50% respectively.
